# Accurate contact predictions for thousands of protein families using PconsC3

**DOI:** 10.1101/079673

**Authors:** Marcin J. Skwark, Mirco Michel, David Menéndez Hurtado, Magnus Ekeberg, Arne Elofsson

## Abstract

Protein structure prediction was for decades one of the grand unsolved challenges in bioinformatics. A few years ago it was shown that by using a maximum entropy approach to describe couplings between columns in a multiple sequence alignment it was possible to significantly increase the accuracy of residue contact predictions. For very large protein families with more than 1000 effective sequences the accuracy is sufficient to produce accurate models of proteins as well as complexes. Today, for about half of all Pfam domain families no structure is known, but unfortunately most of these families have at most a few hundred members, i.e. are too small for existing contact prediction methods. To extend accurate contact predictions to the thousands of smaller protein families we present PconsC3, an improved method for protein contact predictions that can be used for families with as little as 100 effective sequence members. We estimate that PconsC3 provides accurate contact predictions for up to 4646 Pfam domain families. In addition, PconsC3 outperforms previous methods significantly independent on family size, secondary structure content, contact range, or the number of selected contacts. This improvement translates into improved de-novo prediction of three-dimensional structures. PconsC3 is available as a web server and downloadable version at http://c3.pcons.net. The downloadable version is free for all to use and licensed under the GNU General Public License, version 2.

## Introduction

In recent years great progress has been made in the area of residue contact prediction. The vast amount of available sequence data is utilized by direct coupling analysis (DCA) methods to predict contacts between residues with unprecedented quality [1, 2]. This has enabled accurate blind predictions of the structure of soluble proteins [3–5], membrane proteins [6–8], and protein complexes [9, 10]. However, the widespread use of such methods has been limited to protein families with more than 1000 members [11, 12]. Unfortunately, the structure of at least one member of most large families is known (see Fig. S1). This limits the practical usefulness of DCA methods [13] and strongly suggests that methods that accurately predict residue contacts for smaller protein families would be of much greater utility.

Before the advent of DCA methods there has been a longstanding effort in using machine learning techniques to predict residue contacts [14–16]. These methods utilize covariance-based evolutionary information (e.g. mutual information), as well as knowledge based constraints as inputs to a machine learning algoritm. The best non-DCA methods are less dependent on the size of the protein family and although their predictive quality is easily outperformed by DCA on large families, they perform significantly better on smaller families, Figure 1 (a). We have earlier used an iterative machine learning approach, built on the observation that contacts are not randomly distributed, to improve the performance of DCA based contact prediction methods when we developed PconsC2 [17]. Here we propose a way to substantially improve the predictive power, by including state-of-the-art non-DCA predictors among the method’s inputs.

**Fig. 1.**
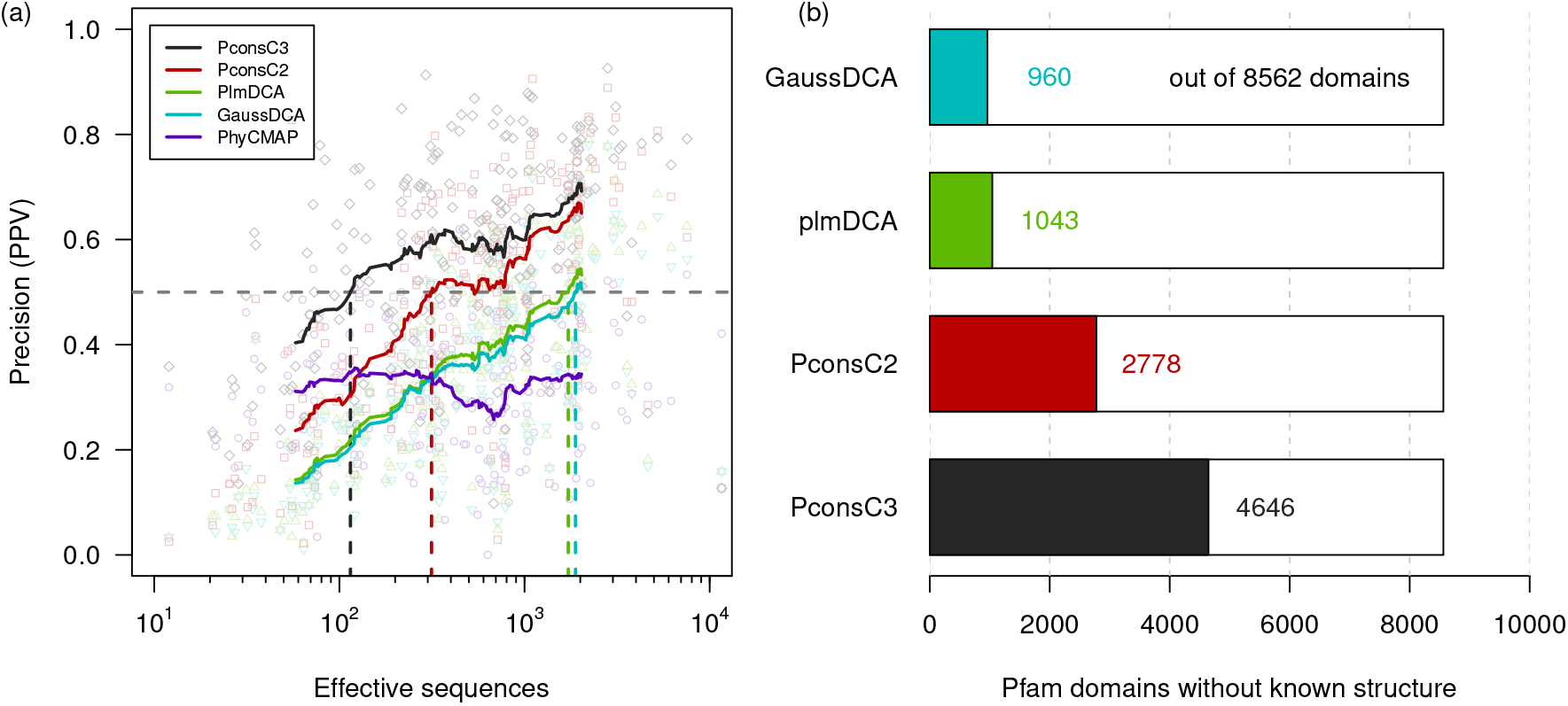
(a) Contact predictor performance on the benchmark dataset measured in Positive Predictive Value (PPV or precision). Performance of the top *N*/2 ranked contacts against protein family size measured in effective sequences, where *N* denotes the number of contacts observed in the native structure. A native contact is defined as a pair of C_*β*_-atoms within a spatial distance of 8Å. The horizontal dashed line marks a precision of 0.5. The vertical lines illustrate at which family size each method reaches this threshold on average. (b) Number of Pfam 29.0 families with unknown structure. A family is defined to have a known structure if there is a significant hit to an entry in PDB that covers more than 75% of the sequence length of the family. In color are shown the numbers of potential target families for each method. These families have at least the number of effective sequences at which the corresponding method reaches an average PPV of 0.5 on our benchmark dataset.

For about half (53%) of the protein families in the Pfam database [18] no structure that covers most of the length can be found in the protein data bank (PDB) [19] (Figure 1b). The distribution of family sizes in the Pfam database shows that the median size of families with known structure (680 effective sequences) is significantly (rank sum p-value *<* 2.2 * 10^−16^) larger than that of families without a known structure (134). The number of potential target families (sufficiently many members, but without known structure) would increase more than three-fold from 1528 to 4973 if accurate predictions could be made from a family with 100 effective sequences instead of 1000 (Supplementary Fig. S1).

PconsC3 combines two DCA methods with contacts predicted using a non-DCA machine learning approach. PconsC3 utilizes the iterative pattern recognition approach introduced in PconsC2 [17]. We find that PconsC3 significantly outperforms earlier methods independent on protein family size, Figure 1 (a), as well as it yields better structural models. The increased accuracy for small proteins leads to a substantial increase in the number of potential targets, from 12% of all Pfam families with unknown structure when using the best DCA method to 54% using PconsC3 instead. These predictions are publically available at http://c3.pcons.net.

#### Significance Statement

One of the longest standing challenges in structural bioinformatics is the protein structure prediction problem, i.e. to predict a protein structure from its sequence. It has recently been shown that, by using accurate residue contact prediction it is possible to predict the structure of many proteins with high accuracy. However, all direct-coupling based contact prediction methods introduced in the last few years require thousands of protein sequences for accurate predictions, but many protein families are smaller. Here, we introduce PconsC3, a method that accurately predicts contacts for families with as few as 100 effective members. When we apply PconsC3 to Pfam we estimate that for 4646 (54%) families of unknown structure we have sufficient coverage for accurate contact predictions.

## 1. Results and Discussion

### Improvement over all protein family sizes

The precision of both DCA methods, plmDCA [20] and GaussDCA [21] as well as that of PconsC2 [17] is strongly dependent on family size. The average Precision (PPV) for PconsC2 for *N*/2 (*N* being the number of native contacts) increases from 0.3 to 0.56 when the average effective family size increases from 100 to 1000 sequences. In contrast, the performance of PhyCMAP [15] is approximately 0.3 for families with between 50 and 2000 effective sequences, Fig. 1 (a). When including PhyCMAP as well as other improvements (see methods) into PconsC3 the performance increases significantly for small families. The average PPV for a 100 effective sequence protein family is 0.47, and increases to 0.60 for a 1000 member family. We have noted that on average a PPV of 0.5 is needed for accurate modeling using the PconsFold [22] pipeline. This average precision is never reached for PhyCMAP, for plmDCA and GaussDCA more than 1700 effective sequences are needed, for PconsC2 314 and for PconsC3 only 115. Even below 100 effective sequences 23% of the benchmark proteins have a PPV larger than 0.5 when using PconsC3.

There are 16295 domains in Pfam 29.0 out of which 8562 do not have a significant match to PDB covering most of the domain length. This means 7733 Pfam domains have at least one representative in PDB and could thus be modeled by homology modelling. If we apply the measurements from Fig. 1 (a) there would be 1043 Pfam domains of unknown structure that could potentially be predicted by the best DCA method. This means these many domains have more than 1700 effective sequences in their alignment. Lowering the threshold of alignment size leads to an increase in the number of potential target domains. With PconsC2 2778 domains could be predicted. This number increases to 4646 when using a method such as PconsC3 that is able to accurately predict contacts from even smaller alignments.

### Performance by type of secondary structure

Figure 2 shows a direct comparison between PconsC3 and other contact predictors. PconsC3 outperforms DCA methods on 207 proteins and PhyCMAP on 195 proteins independent on the type of secondary structure. Compared to PconsC2, PconsC3 performs better in 166 out of 210 proteins of the benchmark dataset. Some of the largest improvements are made for *α **—** β* (cross in Fig. 2) and mainly-*β* proteins (triangle), whereas PconsC2 performs exceptionally well for one short *α*-helical protein (PDB: 1ediA).

**Fig. 2.**
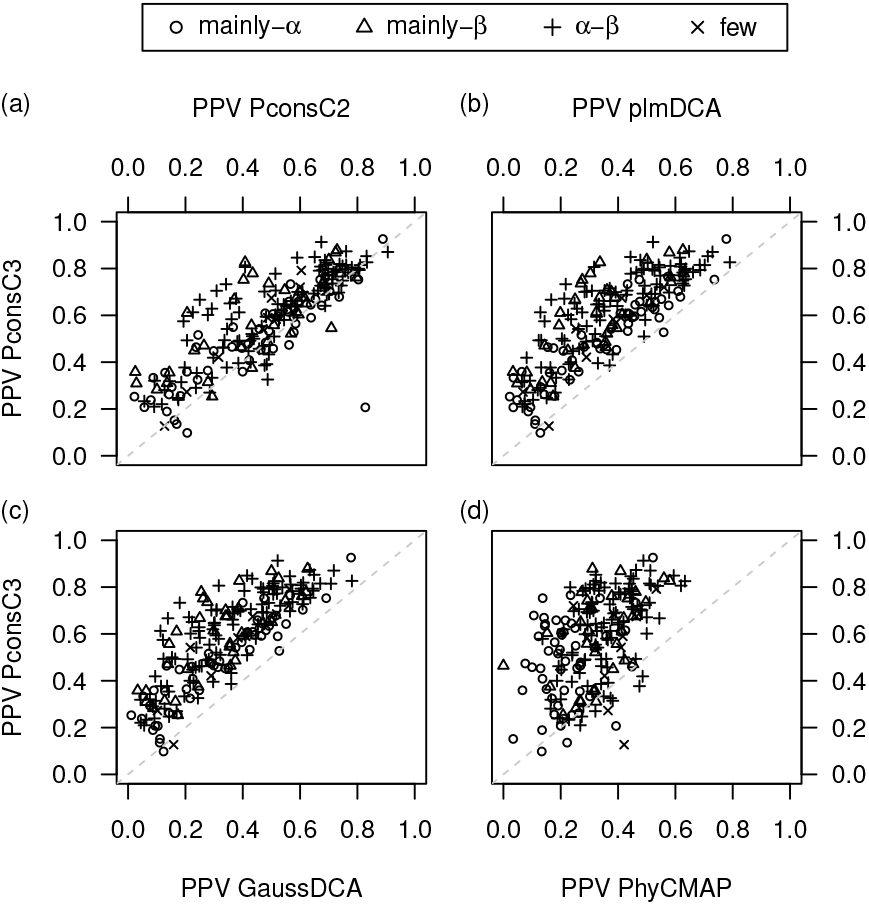
Direct performance comparison between PconsC3 and other methods on the benchmark dataset. Proteins were assigned secondary structural classes based on their ECOD architecture assignment. Symbols represent the class of a protein.

Table 1 shows the performance of contact predictors on different types of secondary structure. The first column lists performance on all proteins of the test set. Overall PconsC3 performs best for all classes independent on rank (Supplementary Fig. S2). On average it predicts more than 57% of *N/2* contacts correctly, compared to 48% for PconsC2, 36% for the best DCA method, and 32% for PhyCMAP. The improvement is largest on mainly-*β* proteins with a 31% increase in PPV of PconsC3 over PconsC2 and 84% over plmDCA. This can be attributed mostly to PhyCMAP performing better than DCA in this particular class. In the *α − β* class PhyCMAP is worse than DCA, while for mainly-*α* proteins both DCA methods clearly outperform PhyCMAP. Within the DCA methods plmDCA always performs better than GaussDCA. Although shown in Fig. 2, the class of few secondary structural elements was omitted in the table due to a small sample size (see Methods).

**Table 1.**
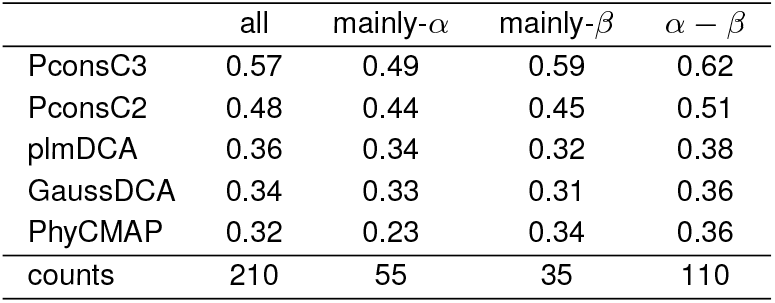
Average PPV of top *N*/2 predicted contacts on the benchmark dataset for different secondary structural classes.

| | all | mainly-α | mainly-β | $\alpha - \beta$ |
| --- | --- | --- | --- | --- |
| PconsC3 | 0.57 | 0.49 | 0.59 | 0.62 |
| PconsC2 | 0.48 | 0.44 | 0.45 | 0.51 |
| plmDCA | 0.36 | 0.34 | 0.32 | 0.38 |
| GaussDCA | 0.34 | 0.33 | 0.31 | 0.36 |
| PhyCMap | 0.32 | 0.23 | 0.34 | 0.36 |
| counts | 210 | 55 | 35 | 110 |

### Predicting long range contacts

structure prediction [23]. There is a striking difference between PhyCMAP and DCA based methods for long-range contacts. PhyCMAP predicts short to medium ranged contacts (with a sequence separation from 5 up to 23 residues) with higher quality than long-range contacts (Fig. 3a). For short-range contacts (up to 12 residues separation) PhyCMAP is actually on par with PconsC2 and significantly better than DCA methods, while it is significantly worse for long-range contacts. Although PconsC3 outperforms DCA methods independently of the sequence separation of contacting residues (Fig. 3a), it is clear that the increase in precision of PconsC3 in relation to PconsC2 and DCA methods is gradually larger for shorter contact ranges, suggesting that it benefits from the good performance of PhyCMAP in this range. On long-range contacts PconsC3 performs best for smaller protein families (Fig. 3b). This and the fact that the gap between PconsC2 and DCA methods decreases for long-range contacts in small families indicate the value of including a non-DCA method to PconsC3.

**Fig. 3.**
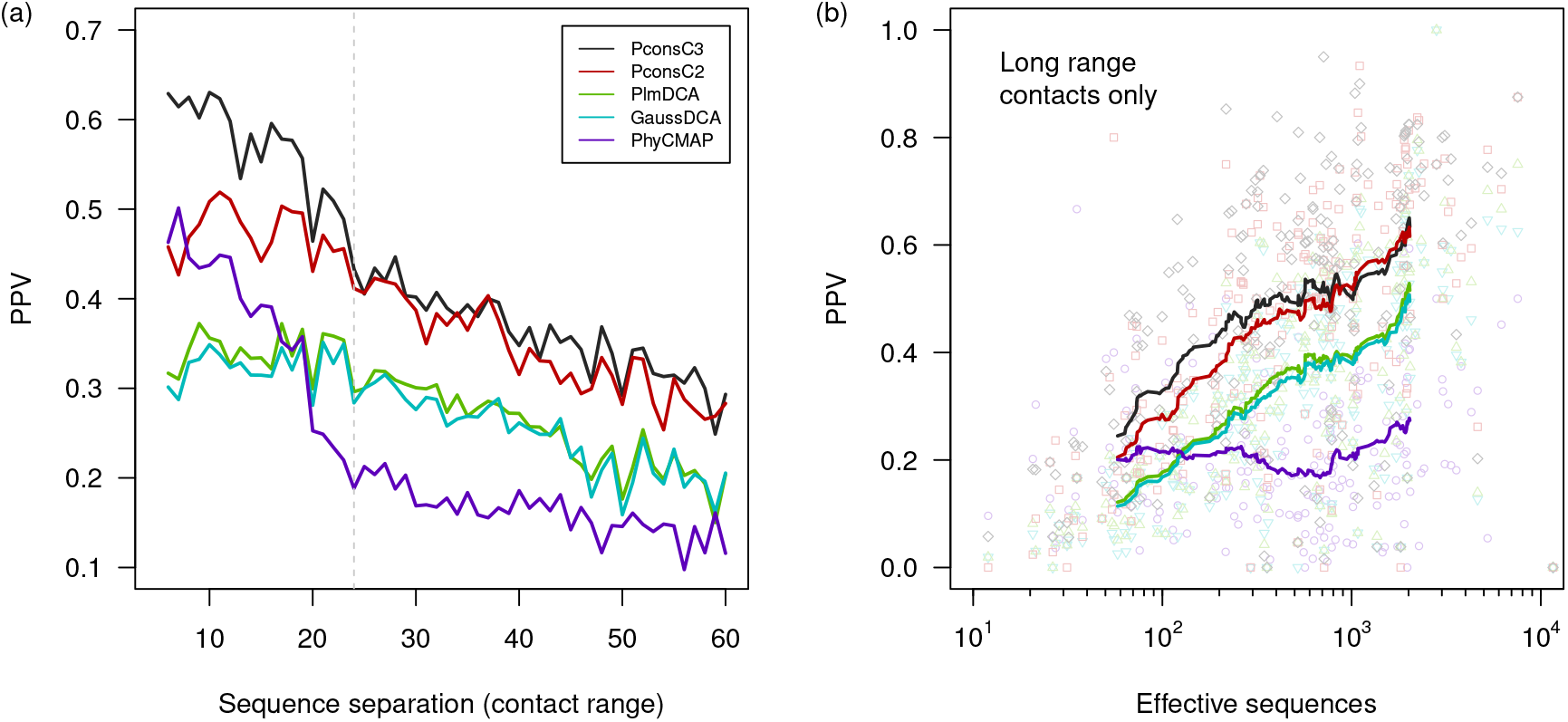
(a) PPV on the top *N*/2 contacts at a specific sequence separation (number of residues between those participating in a contact). A minimum sequence separation of five residues was used to filter out local interactions of neighboring residues or helices. Long-range contacts have a separation of at least 24 residues (everything to the right of the dashed line). (b) Long range contact predictor performance in PPV against protein family size measured in effective sequences.

### Estimation of contact map quality

The average PconsC3 contact scores can be used as a good indicator for contact map quality. Figure 4 (a) shows that the average contact score of the top ranked contacts has a Pearson correlation *r* of 0.61 against PPV. However, we noted the test dataset also includes alleged multi-domain proteins, i.e. proteins where most of the sequences in the alignments does not cover the entire domain, such as 2csmA and 2ejnA, Supplementary Fig. S3 (a and b). Most of the proteins with high average contact score and low PPV fall into that category (gray dots in the lower right region of Figure 4 (a)). This leads to the assumption that PconsC3 is overestimating the predictions in such cases. When ensuring proteins are mostly covered by at least half of all sequences (black dots) *r* increases to 0.83 (*r*_*covered*_) showing that the average PconsC3 score is an excellent estimator of the contact map quality in single domain proteins.

**Fig. 4.**
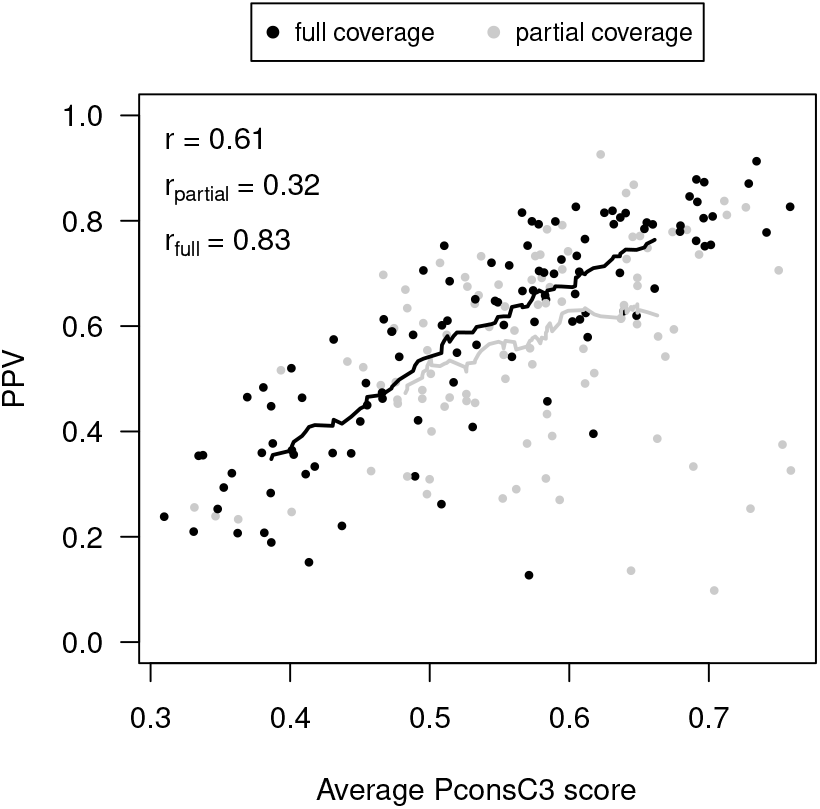
PconsC3 score as an estimator for PPV. Pearson correlation coefficient is denoted *r* and *r*_full_ of all proteins and those with at least 80% of its residues covered by more than 50% of all sequences in the family alignment (length coverage), respectively. The dashed and solid red lines indicate a moving average with window size of 50 for all proteins and those with high length coverage, respectively.

### Structure prediction

The more accurate contact maps of PconsC3 improve structure prediction, confirming earlier observations [22], Fig. 5 (a). The small improvement in average TM-score [24] when using PconsC3 contacts is significant (t-test p-value < 1.5 · 10^−2^) and independent of the family size of the target protein. However, there are still many proteins with large families and supposedly good contact maps, for which PconsFold fails to converge properly (Supplementary Fig. S4). Using a more elaborate folding protocol [25] or generating more than 2000 Rosetta decoys might improve this situation. Anyhow, when using predicted contacts from PconsC3 the number of proteins with a TM-score of 0.5 or higher, meaning the fold has most likely been correctly identified, increases from 55 to 75.

**Fig. 5.**
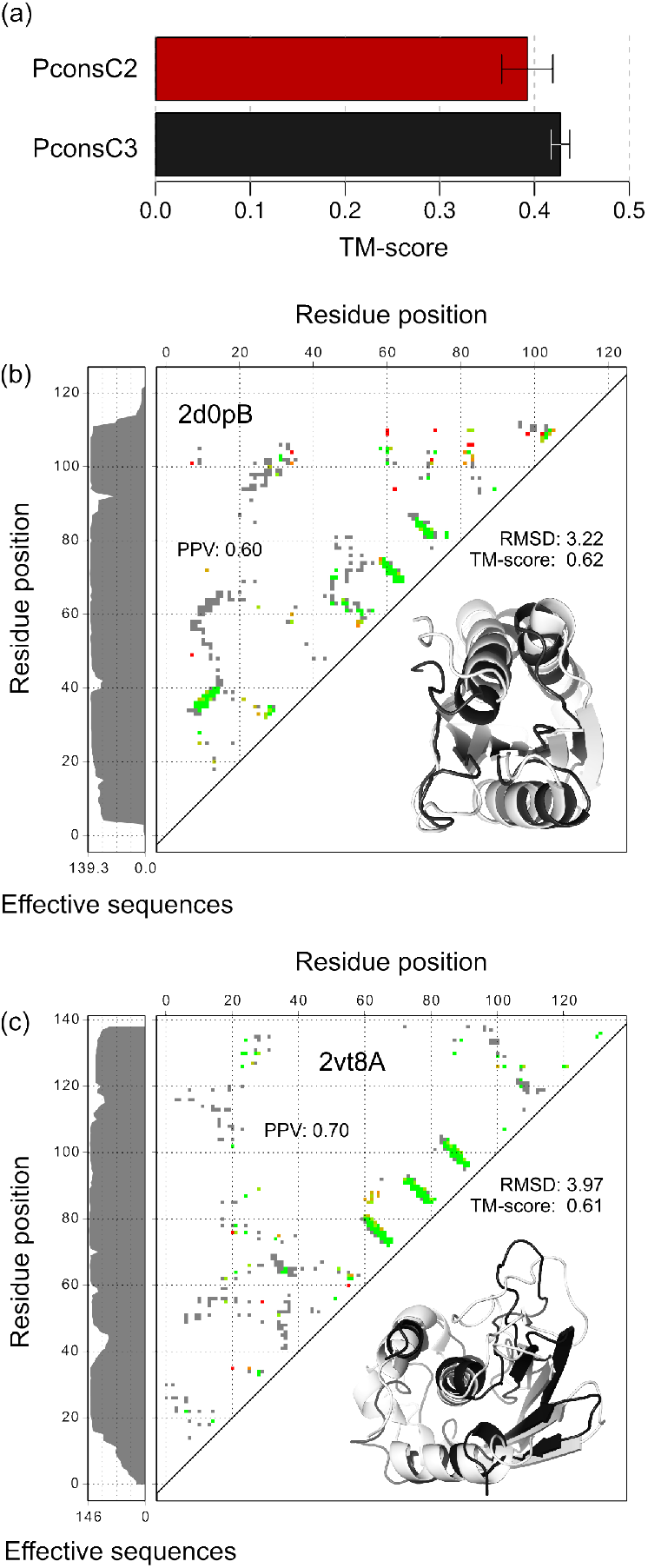
Structure prediction. (a) TM-score of PconsFold when using PconsC2 (red) or PconsC3 (black) contact predictions. (b) Contact map for Diol dehydratase reactivase ATPase-like domain (Pfam: PF08841, PDB: 2d0pB) in the upper left triangle. Grey dots indicate native contacts in the PDB structure, green are correctly predicted contacts while yellow to red are false positive predictions. Left the sequence coverage measured in effective sequences. The lower right triangle shows the structure predicted with PconsFold using PconsC3 contacts (black) superimposed onto the native structure from PDB (light gray) (c) Contact map and structure for the PI31 proteasome regulator N-terminal (PF11566, 2vt8A).

The main advantage of PconsC3 over PconsC2 and DCA methods is that it can accurately predict the contacts for smaller protein families. The Diol dehydratase reactivase ATPase-like domain (PF08841) only contains 139 effective sequences but both the contact map and the model are in excellent agreement with the native structure (2d0pB), Fig. 5(b). The TM-score of the model is 0.61 while a model based on PconsC2 only has a TM-score of 0.40. The PI31 proteasome regulator N-terminal (PF11566) has 146 effective sequences and for this protein a TM-score of 0.61 is reached, Fig. 5(c).

### Blind prediction of T0872 in CASP12

The submission phase for the twelfth Critical Assessment of Techniques for Protein Structure Prediction (CASP12) recently finished and the official evaluations are running as of writing this manuscript. Initial evaluation of CASP12 targets that can already be found in PDB revealed that the combination of PconsC3 and PconsFold successfully predicted the structure of Target T0872. Model 3 (Pcons-net_TS3) has the highest TM-score of 0.74 followed by Pcons-net_TS1. All five Pcons-net submissions rank among the top 10 models for this target. Figure 6 shows the predicted contact map as well as the structure of Model 3, both overlaid on the native contacts (gray) and structure (white), respectively. The high precision of the predicted contacts can be attributed to the large sequence coverage of 3817 effective sequences in the alignment. Furthermore, PconsFold was able to converge to a near native structure most likely due to the small size of the protein. For some larger targets this was not the case.

**Fig. 6.**
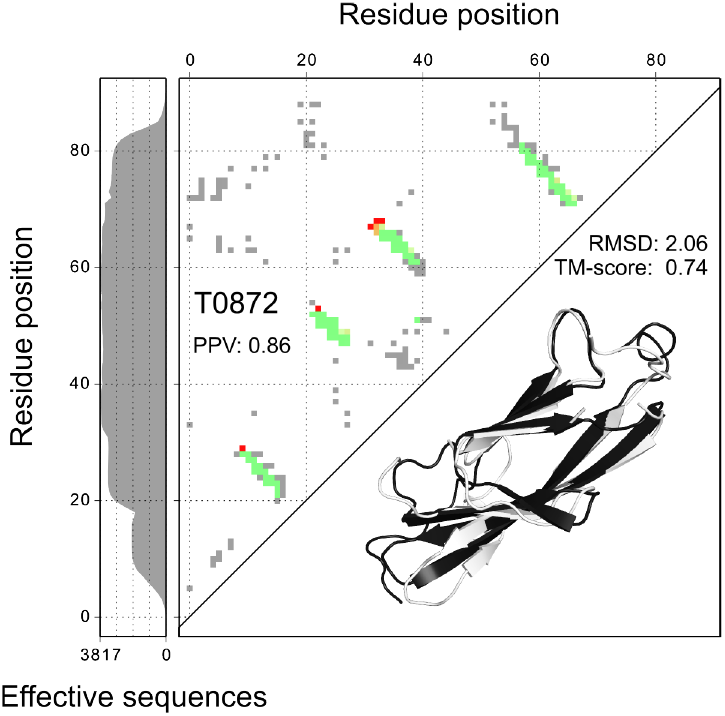
Blind prediction of target T0872 in CASP12. Predicted contact map on top of the native in the upper left triangle. The lower right triangle shows the structure predicted with PconsFold using PconsC3 contacts (black) superimposed onto the native structure from PDB (light gray).

## Materials and Methods

### Datasets

PconsC3 has been trained on a set of 180 protein families (supplementary Table S1). This training set comprises of 150 protein families from the original PSICOV dataset [26] plus 30 additional families with a small number of members from the test set as described in [17].

All evaluation has been made on a dataset of 210 (supplementary Table S2) proteins without any homology to any protein in the training set. All PDB IDs were matched against the ECOD [27] domain assignment from 2016-03-28. This set was obtained from the set used in the development of PconsC2 [17] and homology reduced such that no protein included in the test set shared an ECOD H-class with any of the proteins in the new training dataset. This homology reduction is much more stringent than using sequence information alone. The final list of proteins used as well as their ECOD H-class number and number of effective sequences are found in Tables S1 and S2. Alignments were created using HHblits [28] version 2.0.15 on the uniprot20 database bundled with HHsuite (date: 2016-02-26) with an e-value of 1. In order for HHblits to output and align all sequences the parameter `-all` has been used and `-maxfilt` and `-realign_max` were both set to 999999. These alignments were used as input for the DCA methods.

For the evaluation of Pfam domains the HHsuite database of Pfam 29.0 (date: 2016-05-03) was used to scan Uniprot at an e-value threshold of 1 using HHblits. The resulting set of alignments was then analyzed for effective number of sequences. For each domain the sequence that was highest ranked by HHblits has been defined as the domain representative. The length of a domain has been set to the length of its representative sequence. HHsearch version 3.0.0 was used to scan each family against the HHsuite database of protein data bank (PDB) sequences (date: 2016-03-02) to determine whether a given Pfam family has known or unknown structure. A hit has been considered significant if its E-value was below 10^−3^ and if it covered at least 75% of the length of the family.

### Secondary structural classes

To classify the dataset into the secondary structural classes mainly-*α*, mainly-*β*, and *α − β,* we used the architecture assignment of ECOD for the PDB IDs of our benchmark set. ECOD uses a scheme with seven structural classes that we mapped into three in order to increase sample size and thus statistical significance of each class. The following mapping was applied: *α/β, α − β* and *α*+*β* to *α − β*; *α to* mainly-*α*; *β* and *extended* to mainly-*β*. The secondary structural class *few* was omitted as it only contained 10 proteins. Supplement Figure S3 shows a table analogous to 1, but with the original ECOD classification (including *few).*

### Contact prediction

Julia implementations have been used for both plmDCA and GaussDCA, which are available on GitHub at https://github.com/pagnani/plmDCA and https://github.com/carlobaldassi/GaussDCA.jl, respectively. Both require Julia 0.3 or higher. PhyCMAP was obtained at http://raptorx.uchicago.edu/download/. Regularization strength of plmDCA was set to 0.02. GaussDCA and PhyCMAP were run with default parameters. The DCA methods were directly run on the alignments described above, whereas PhyCMAP runs its own workflow and thus uses its own alignment as described in [15]. PconsC2 was run as described before [17].

### PconsC3

Figure S6 illustrates the workflow of PconsC3. Input features comprise contact predictions by plmDCA, GaussDCA as well as PhyCMAP, secondary structure prediction by PSIPRED 3.0 [29], and solvent accessibility prediction by NetSurfP 1.1 [30]. In PconsC3 PhyCMAP can be replaced by another contact predictor and we have successfully used CMapPro [16] with similar accuracy, data not shown.

Additionally, CD-HIT is run to generate statistics about the alignment (i.e. alignment depth at different sequence similarity cut-offs). The initial layer of PconsC3 takes these features as input and uses a random forest to predict a score for each possible contact. In contrast to previous work, PconsC3 applies pattern recognition already in the first layer. This results in an intermediate contact map. Every following layer uses all the initial features plus the output from the previous layer, given as a window of 11 by 11 residues around the current contact. Note that the pattern recognition method used in PconsC3 is analogous to convolutional layers of deep learning, as described in detail earlier [17].

The initial layer of PconsC3 (PconsC3-l0) shows an increased precision over PconsC2 independent of the number of top-ranked predicted contacts used for evaluation (Fig. S2). The precision increases for each layer to saturate at the third layer. In contrast to PconsC2 the fourth and fifth layers does not increase the performance.

Each of the Random Forests comprising PconsC3 consists of 100 trees trained based on optimization of Gini impurity, with a constraint on node split with at least 100 samples per leaf. To reduce the memory footprint of training, as well as to prevent overfitting, starting from layer 1, we have disregarded a randomly chosen subset of 30% of the training samples, which appears to improve the generalizability of resulting statistical models.

At https://github.com/mskwark/PconsC3/ instructions on how to setup and run PconsC3 locally are found. PconsC3 can also be used from a web-server at http://c3.pcons.net/, where predictions for ≈ 14000 of the sufficiently large Pfam domain families also can be found.

### Selecting top ranked contacts

We analyzed the top ranked contacts using half of the number of observed contacts (*N*/2, dashed vertical line in Figure S2). This number roughly corresponds to the length of the protein (*L*) (Supplementary Fig. S7), i.e. the same number of contacts used to analyze precision (PPV) earlier [17].

A native contact between two residues is present if their *C*_*β*_-atoms is within 8Å. The contact score was used to rank predicted contacts and the top *N*/2 contacts were used for evaluation. This allows for a fair comparison between the methods, while being easy to interpret, e.g. if a method has a PPV of 0.5 at *N*/2 contacts, one can say that this method correctly predicts 25% of all observed contacts. Thereby, false and true negatives are implicitly taken into account. For this reason we decided to choose a cut-off based on *N* instead of the widely used cut-off based on the length of the input sequence *L.*

### Metrics

Effective sequences is defined in analogy to [31] as:

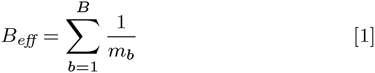

where *m*_*b*_ is the number of sequences with at least 90% sequence identity 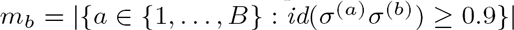.

The quality of a predicted contact map is measured in positive predictive value (PPV), or precision:

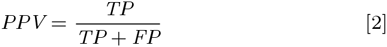

where TP is the number of predicted contacts that match a contact in the native structure (true positives) and FP the number of predicted contacts that don’t (false positives).

TM-score [24] is used to measure the similarity between predicted and native structure. To enable fair comparison with PconsC2, we used the same cut-off (top 1.5 *·L* contacts, where L denotes sequence length) as in [17] to select contacts for PconsFold. Preliminary observations clearly indicate that better performance can be obtained using another scheme, but we have not systematically evaluated this.

## ACKNOWLEDGMENTS

This work was supported by grants from the Swedish Research Council (VR-NT 2012-5046) to A.E.) and Swedish e-Science Research Center (SeRC). Computational resources at the National Supercomputing Center were provided by SNIC. The website is maintained by the Bioinformatics Infrastructure for Life Sciences (BILS). The authors thank Nanjiang Shu for setting up the website. The authors declare that they have no competing financial interests.

## Supplementary Figures

**Fig. S1.**
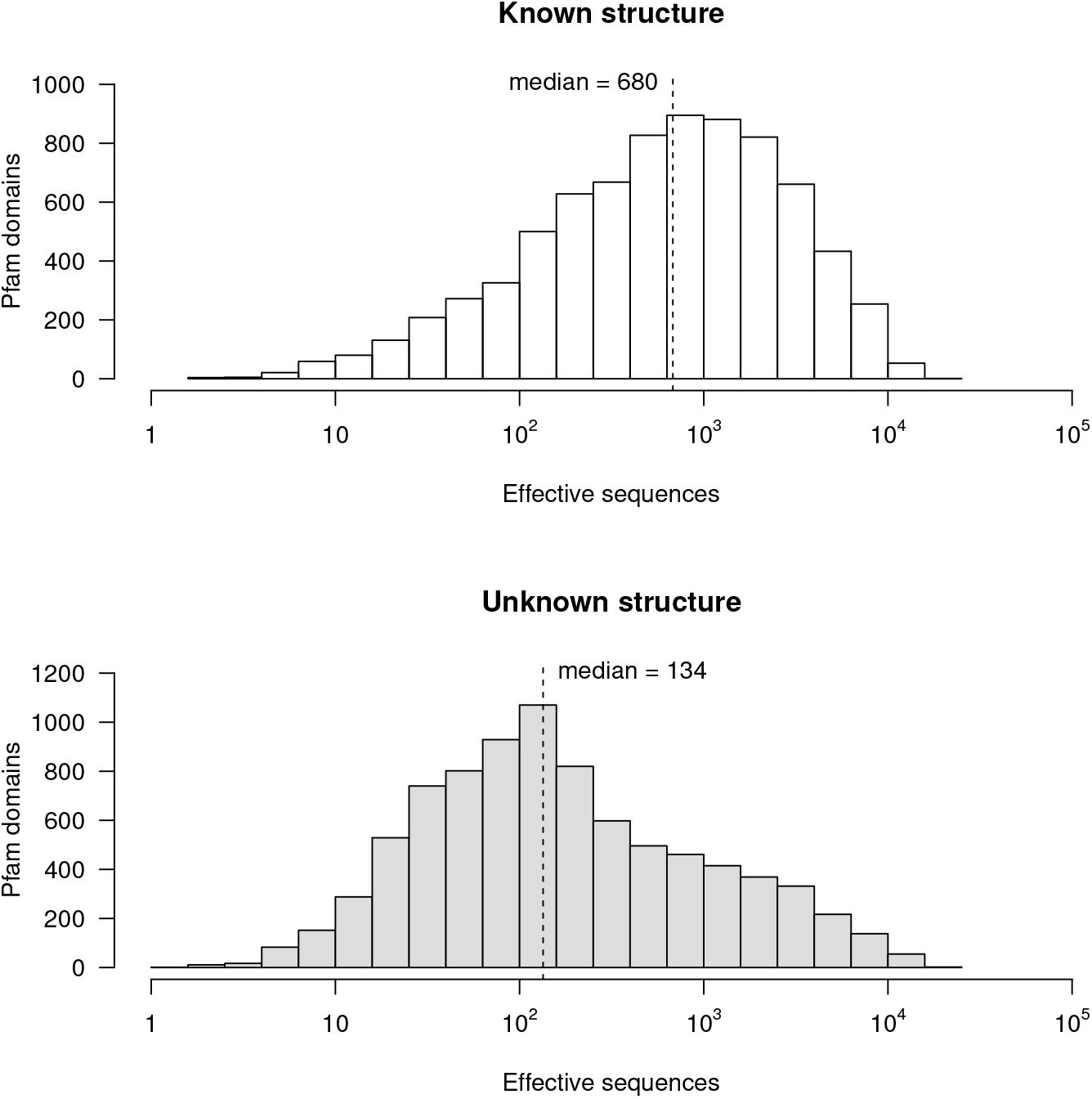
Pfam family sizes measured in effective sequences. (a) Histogram over the size of all families with at least one hit in PDB that covers more than 80% and (b) of all families without such hit. (c) Cumulative counts of Pfam families over their size measured in effective sequences of the underlying alignment. There are 16295 protein families in Pfam 29.0 out of which 8562 do not have a significant hit to a structure in PDB, which would cover more than 75% of their length. The vertical line segments represent 100 and 1000 effective sequences. Horizontal line segments connect these to the corresponding number of Pfam families.

**Fig. S2.**
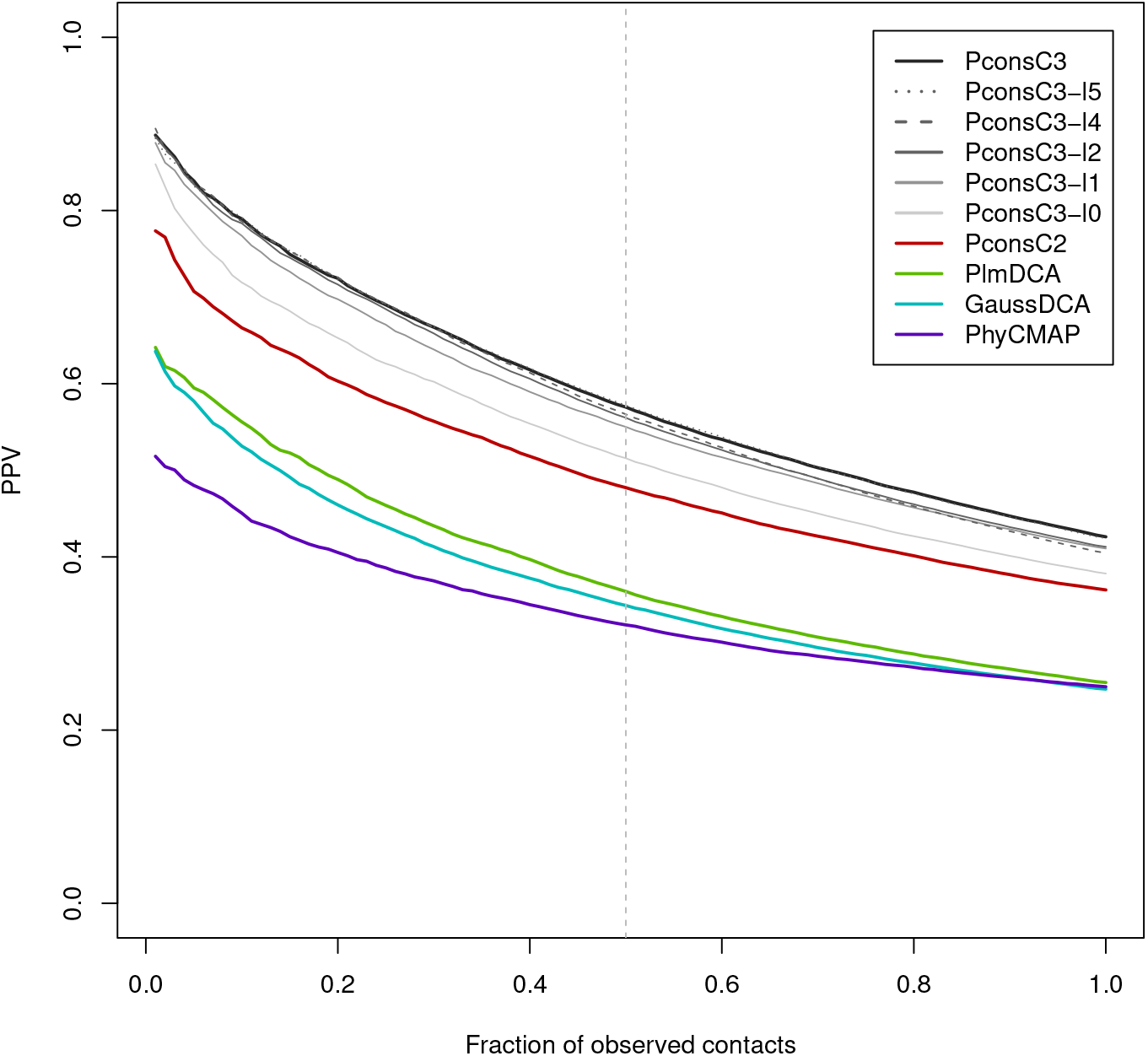
Contact predictor performance on the independent test set measured in Positive Predictive Value (PPV or precision). Performance against the number of top ranked predicted contacts measured as a fraction of contacts observed in the native structure *N*. The dashed vertical line indicates the number of contacts used further on.

**Fig. S3.**
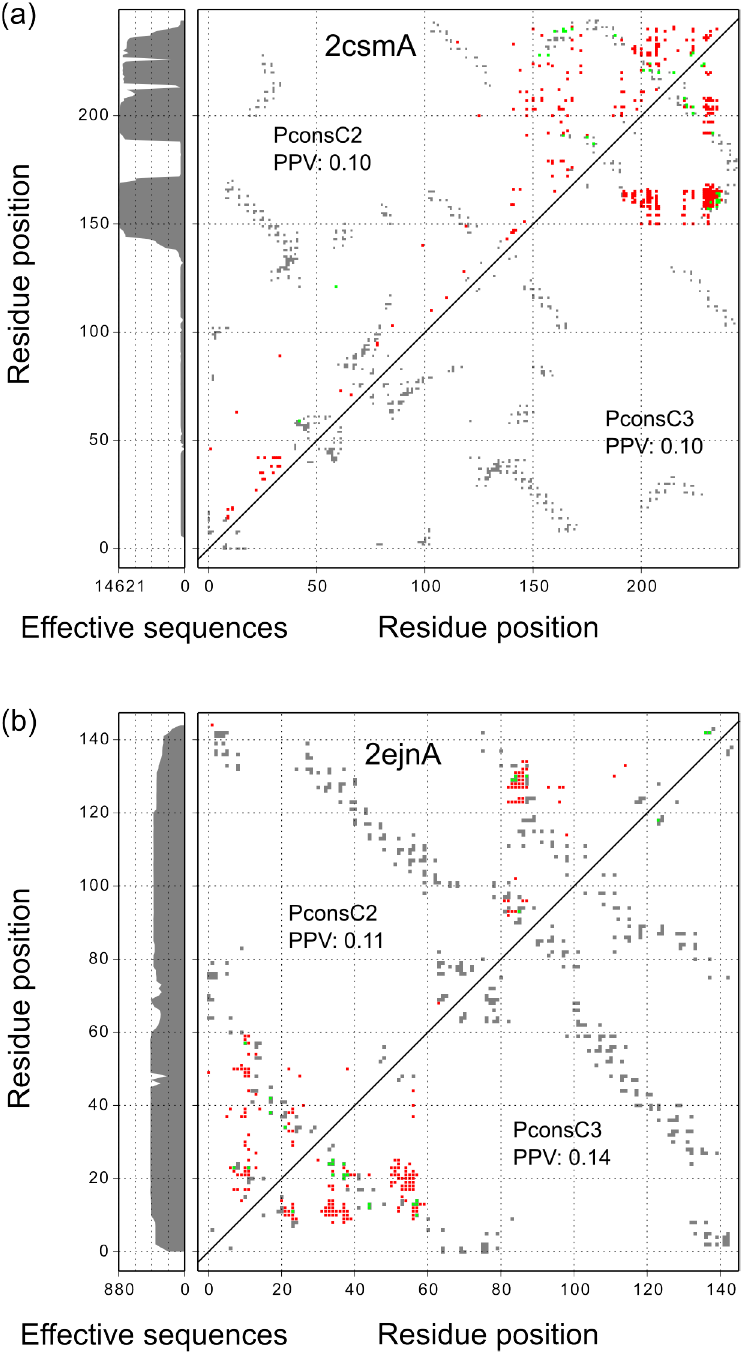
Example proteins where most of the aligned sequences do not cover the entire length. The alignment coverage is indicated by the vertical panel on the left-hand side, where the width of the gray area represents the number of effective sequences at that position. (a) PDB: 2csmA is only covered in the terminal region (b) PDB: 2ejnA is mostly covered by only half of the sequences, thus consists of two separate parts.

**Fig. S4.**
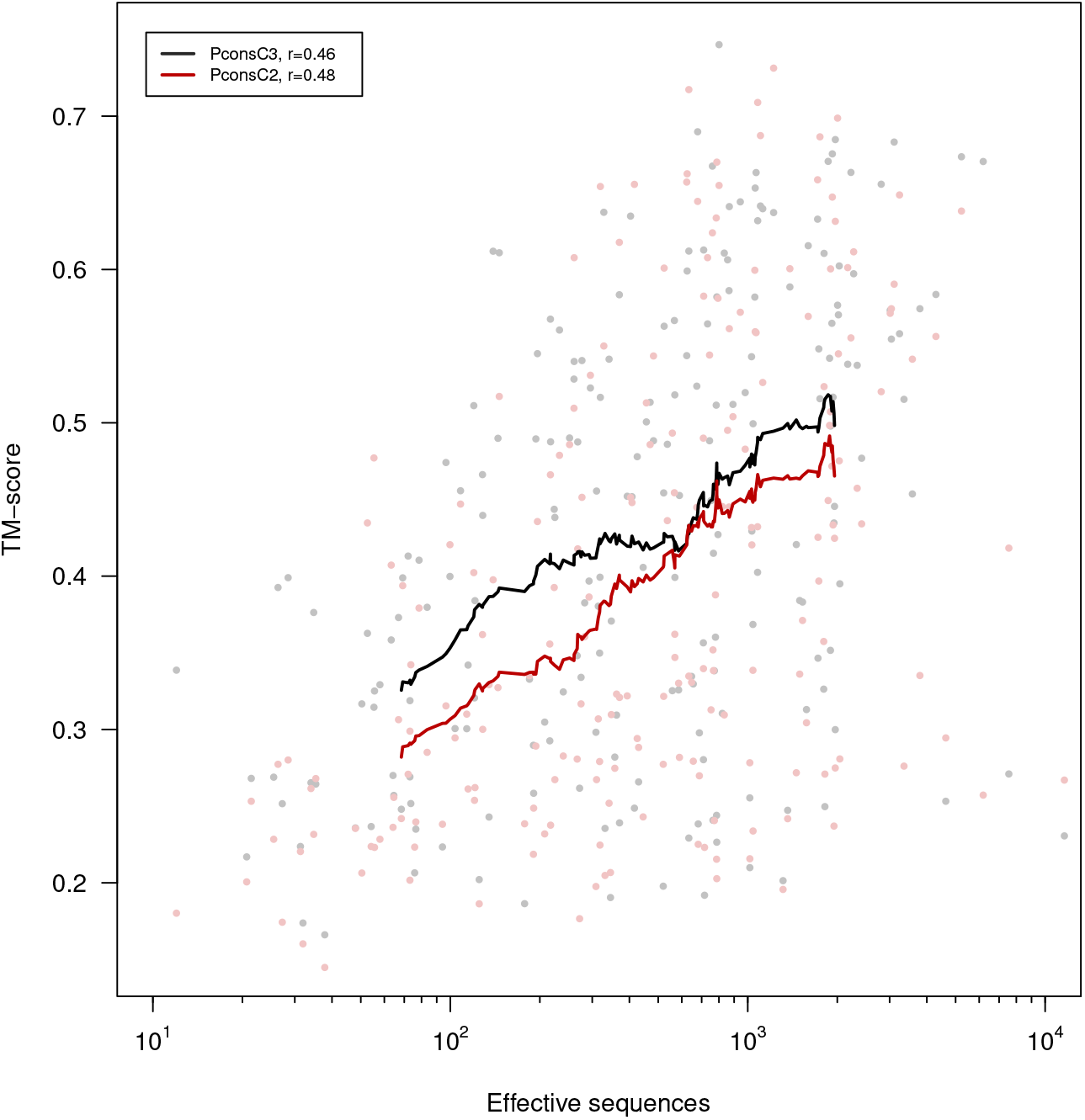
TM-score of PconsFold against family size when using PconsC3 (black) or PconsC2 (red) contact predictions. Pearson correlation coefficient is denoted *r* and the lines indicate moving average with a window size of 50.

**Fig. S5.**
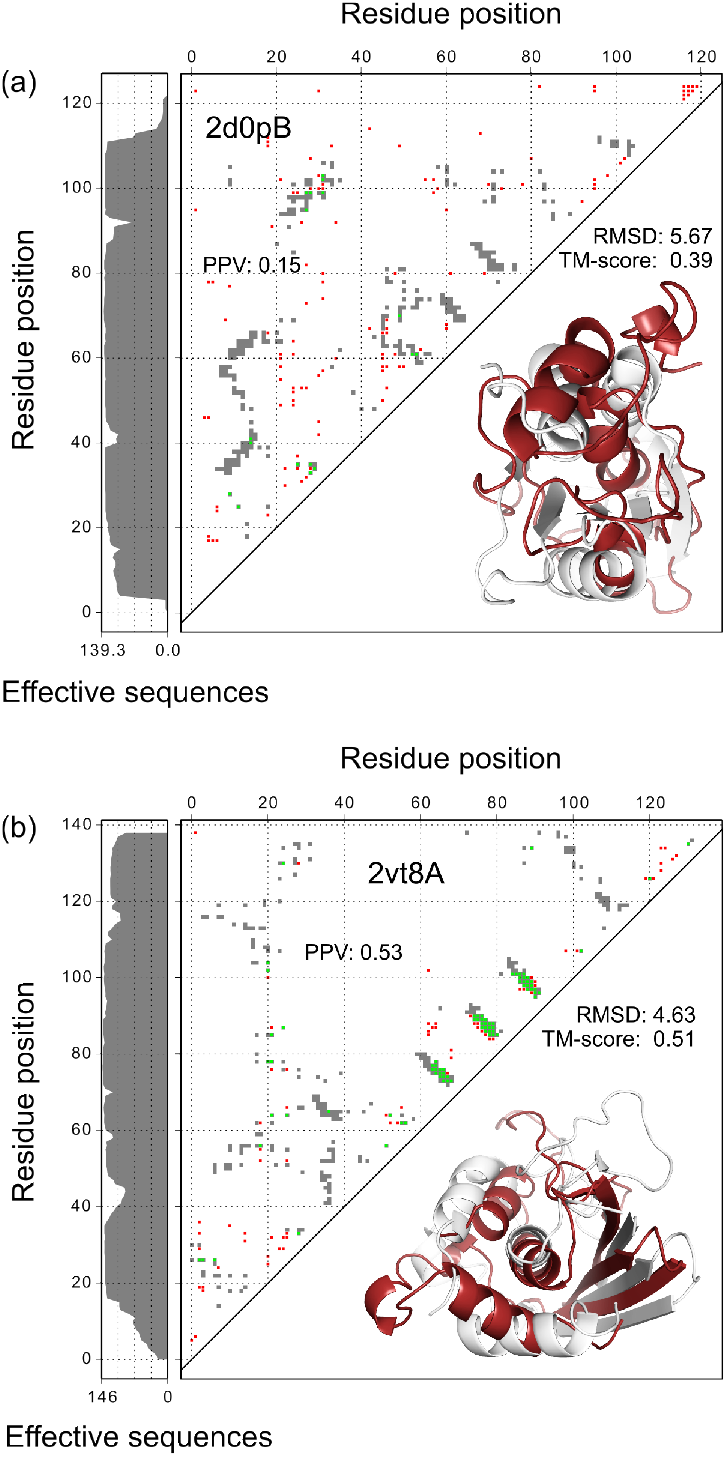
Structure prediction. (a) Contact map for Diol dehydratase reactivase ATPase-like domain (Pfam: PF08841, PDB: 2d0pB) in the upper left triangle. The lower right triangle shows the structure predicted with PconsFold using PconsC3 contacts (black) superimposed onto the native structure from PDB (light gray) (b) Contact map and structure for the PI31 proteasome regulator N-terminal (PF11566, 2vt8A).

**Fig. S6.**
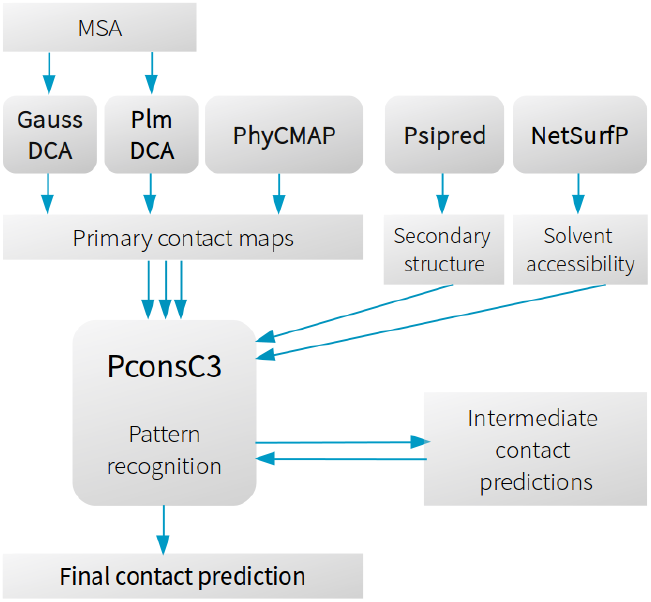
PconsC3 workflow. GaussDCA and PlmDCA are combined with the non-DCA method PhyCMAP and additional secondary structure and solvent accessibility features. PconsC3 combines all features and iteratively predicts intermediate contact maps. In every iteration predictions from the previous layer are used as additional input providing a description of the contact pattern.

**Fig. S7.**
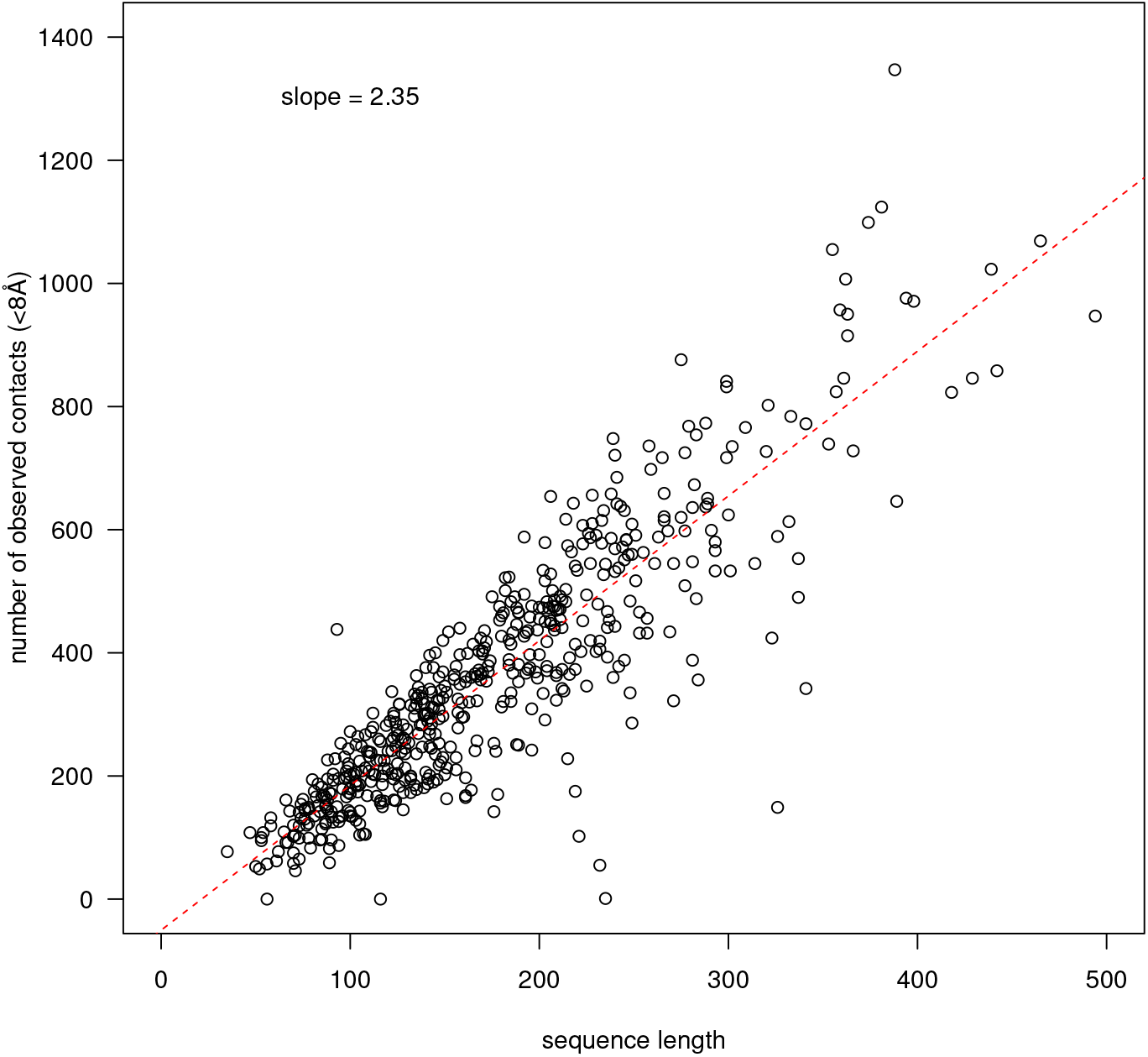
Sequence length against the number of contacts observed in the native structure (*N*) for all proteins in both training and test set. Native contacts are defined as C_*β*_-atoms closer than 8Å. Counted are all native contacts with a sequence separation above 5 residues.

---

**Table S1.**
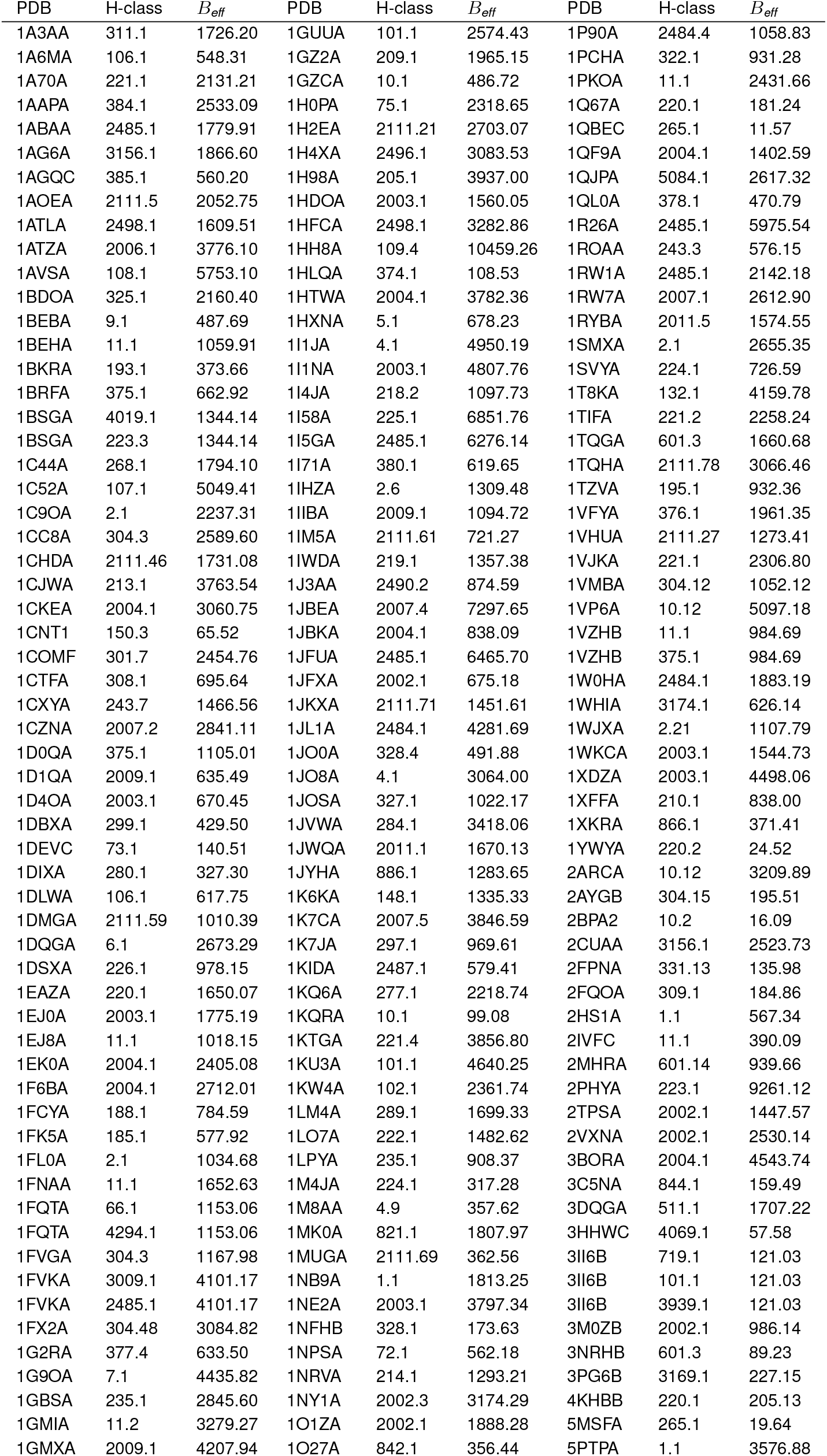
Training dataset

**Table S2.**
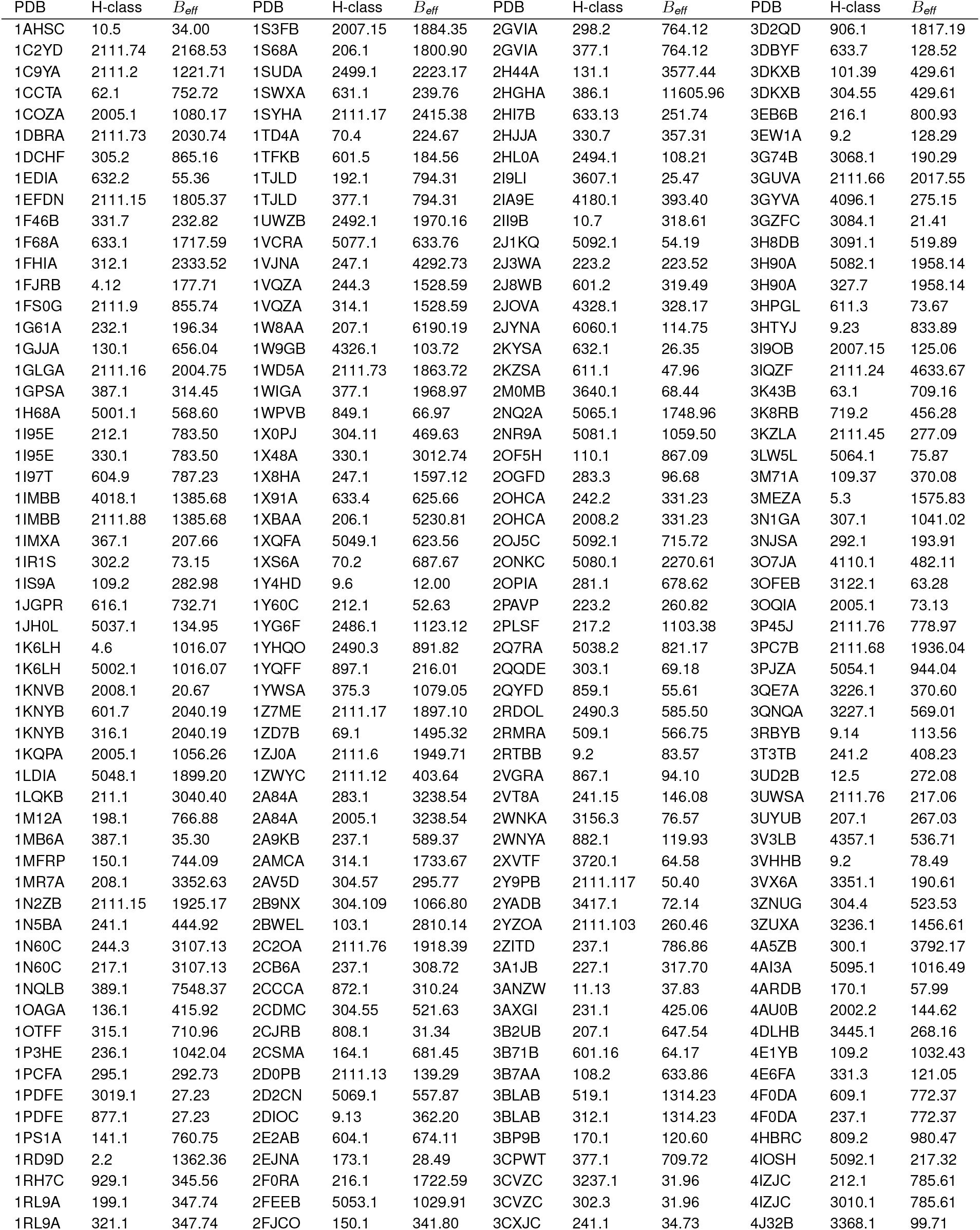
Test dataset

**Table S3.** Average PPV of top *N*/2 predicted contacts on the independent test set for the original ECOD secondary structure classes.

| method | all | $\alpha/\beta$ | $\alpha+\beta$ | $\alpha$ | $\beta$ | extended | few | mixed |
| --- | --- | --- | --- | --- | --- | --- | --- | --- |
| PconsC3 | 0.57 | 0.62 | 0.61 | 0.49 | 0.59 | 0.46 | 0.54 | 0.77 |
| PconsC2 | 0.48 | 0.55 | 0.49 | 0.44 | 0.46 | 0.39 | 0.44 | 0.78 |
| PlmDCA | 0.36 | 0.43 | 0.35 | 0.34 | 0.32 | 0.36 | 0.37 | 0.61 |
| GaussDCA | 0.34 | 0.40 | 0.33 | 0.33 | 0.31 | 0.36 | 0.36 | 0.60 |
| PhyCMAP | 0.32 | 0.34 | 0.37 | 0.23 | 0.35 | 0.00 | 0.39 | 0.45 |
| counts | 210 | 40 | 69 | 55 | 34 | 1 | 10 | 1 |

